# The coupling of global brain activity and cerebrospinal fluid inflow is correlated with Alzheimer’s disease related pathology

**DOI:** 10.1101/2020.06.04.134726

**Authors:** Feng Han, Jing Chen, Aaron Belkin-Rosen, Yameng Gu, Liying Luo, Orfeu M. Buxton, Xiao Liu, Alzheimer’s Disease Neuroimaging Initiative

**Affiliations:** Department of Biomedical Engineering, The Pennsylvania State University, PA, USA; Department of Sociology & Criminology, The Pennsylvania State University, PA, USA; Population Research Institute, The Pennsylvania State University, PA, USA; Department of Biobehavioral Health, The Pennsylvania State University, PA, USA; Institute for Computational and Data Sciences, The Pennsylvania State University, PA, USA

## Abstract

The glymphatic system plays an important role in clearing the amyloid-β and tau proteins that are closely linked to Alzheimer’s disease (AD) pathology. Glymphatic clearance, as well as amyloid-β accumulation, is highly dependent on sleep, but the sleep-dependent driving forces behind cerebrospinal fluid (CSF) movements essential to the glymphatic flux remain largely unclear. Recent studies have reported that widespread, high-amplitude spontaneous brain activations in the drowsy state and during sleep, which are shown as large global signal peaks in resting-state fMRI, is coupled with the CSF movements, suggesting their potential link to the glymphatic flux and metabolite clearance. By analyzing multimodal data from the Alzheimer’s Disease Neuroimaging Initiative project, here we showed that the coupling between the global fMRI signal and CSF influx is correlated with AD-related pathology, including various risk factors for AD, the severity of AD-related diseases, the cortical amyloid-β level, and the cognitive decline over a two-year follow-up. These results provide critical initial evidence for involvement of sleep-dependent global brain activity, as well as the associated physiological modulations, in the clearance of AD-related brain waste.

## Introduction

The pathogenesis of Alzheimer’s disease (AD) is widely believed to be driven by the aggregation of toxic proteins, e.g., the amyloid-β (Aβ) and tau, that cannot be adequately cleared from the brain^1–3^. The “glymphatic system” plays an important role in the clearance of the toxic proteins in the extracellular interstitial space^4–8^. In this clearance pathway, the movement of the cerebrospinal fluid (CSF) from the periarterial space into the interstitial space, which is facilitated by astroglial aquaporin-4 (AQP4) channels, drives convective interstitial fluid (ISF) flux and interstitial solutes, including Aβ and tau, into the perivenous space surrounding deepdraining veins, from where the effluxed wastes can then be absorbed into the cervical lymphatic system^7,8^. An intriguing finding regarding this glymphatic pathway is that the CSF influx can increase 20 times during sleep compared with the awake condition^4^. This highly sleep-dependent clearance may contribute to the diurnal fluctuation of interstitial Aβ observed in both human and mice^9,10^, as well as the link between AD pathology and sleep^11–13^. However, this strong sleep dependency obscured our understanding of the driving forces behind the CSF influx of the glymphatic process. Arterial pulsation^6,14–16^ and respiration^17,18^, which have been hypothesized to be the major drivers of the glymphatic CSF movements, would not explain this sleep-dependent enhancement because they are expected to remain similar or even decrease during sleep due to reduced physical activities^4^.

The glymphatic process may be related to sleep-dependent global brain activity recently observed with resting-state functional magnetic resonance imaging (rsfMRI)^19–21^. Correlations of rsfMRI blood-oxygen-level-dependent (BOLD) signals have been widely used to measure functional brain connectivity ^22,23^, but the whole-brain mean of resting-state BOLD signals, i.e., the global BOLD signal, had been often regarded as artifact and removed from connectivity analysis^24–26^. Converging evidence, however, suggested that the global BOLD signal is of neural origin^27^, highly dependent on brain state^27,28^, and especially strong during light sleep^29,30^. Consistent with these observations, caffeine^31^ can effectively suppress this global component whereas hypnotic drugs, i.e., midazolam and zolpidem^32–34^, and sleep deprivation^35^ had opposite effects. A recent study combining multimodal imaging data from monkeys and humans^21^ suggested that the global BOLD signal is tightly linked to a specific electrophysiological event of 10-20 seconds duration that occurs during the drowsy state or sleep^20^. This sequential spectral transient (SST) event involves widespread cortical fMRI co-activations and very specific subcortical de-activation in the arousal-promoting regions at the basal forebrain and dorsal midline thalamus, and also shows a characteristic time-frequency signature indicative of transient arousal modulation^20,21^. The large global BOLD changes are also accompanied by strong cardiac^36^ and respiratory^37,38^ modulations. More importantly, the large global BOLD signal fluctuation was recently found to be accompanied by CSF inflows during sleep^39^, suggesting its potential role in the glymphatic process that may be important for AD pathogenesis. Based on all these findings, we hypothesize that the coupling between the global BOLD signal and CSF inflow is related to AD-related pathology.

To test this hypothesis, we examine multimodal data from the Alzheimer’s Disease Neuroimaging Initiative (ADNI)^3^ with a focus on the coupling of the global BOLD signal and CSF signal and its relationship with AD-related neurobiological and neuropsychological measures. We found a strong coupling between the global brain signal and CSF signal, which is significantly attenuated in the AD-related disease groups and correlated with multiple AD risk factors. The BOLD-CSF coupling strength is significantly correlated with the cortical Aβ level and also predicts the cognitive decline in the subsequent two years. The findings suggest an important role of the neural and physiological processes associated with the global brain BOLD signal in the AD pathogenesis.

## Results

### Sample characteristics

We used imaging and behavioral data from 118 subjects in the ADNI project. The sample included 7 AD patients, 62 mild cognitive impairment (MCI) subjects, 18 significant memory concern subjects (SMC) and 31 healthy controls (HC) (see **Table 1** for detailed sample characteristics). They were selected based on the availability of rsfMRI, florbetapir positron emission tomography (PET) measurement of Aβ, and behavioral measures through Mini–Mental State Examination (MMSE). The 118 subjects underwent 158 rsfMRI sessions that are associated with the baseline and follow-up (after ~2 years) Aβ-PET and MMSE measurements (see **Methods** for details about subject selection). This sample allowed us to link the baseline rsfMRI data to longitudinal changes of AD-related markers. Data from 29 out of 118 subjects were collected in three consecutive time points (i.e., three Aβ-PET and MMSE sessions at year 0, 2 and 4, and corresponding rsfMRI sessions at year 0&2), and were then split into two pairs of measurements by re-using the middle time point as the baseline of the third time point. Similarly, we split the data of four subjects with four consecutive measurements and one subject with five consecutive measurements to increase the pairs of measures with a 2-year interval.

**Table 1.**
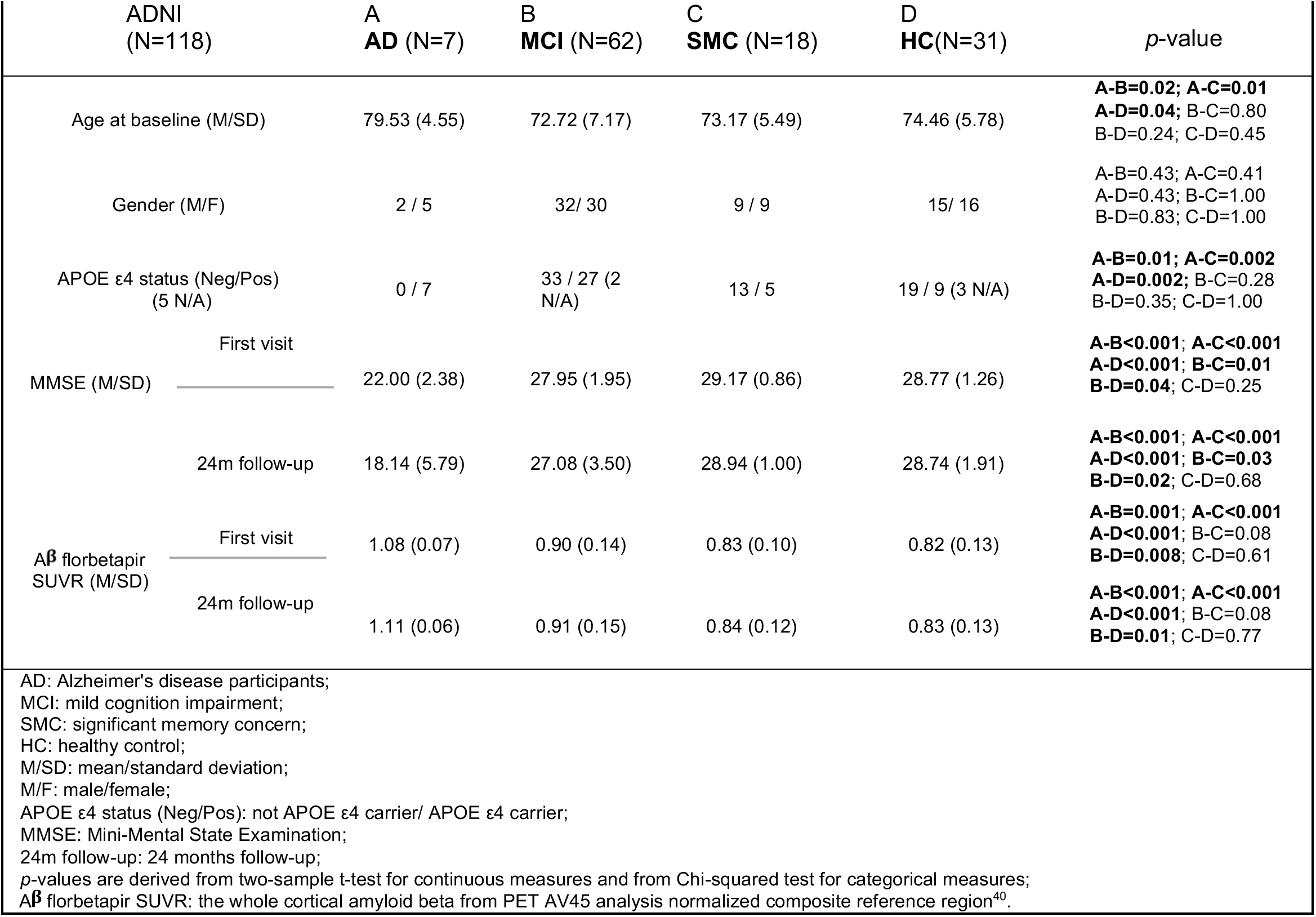
Subject characteristics.

### A coupling between the global BOLD signal and CSF signal

We first examined whether the BOLD-CSF coupling described in the previous study^39^ is also present in the ADNI rsfMRI data collected in relatively old subjects and with lower spatial and temporal resolutions. Similar to the previous study, we extracted CSF signals from the bottom slice of rsfMRI acquisition, which is around the bottom of the cerebellum, to maximize sensitivity to the CSF inflow effect^39^. The CSF regions can be easily identified in the T2*-weighted fMRI images since they appear much brighter than surrounding tissues (the right panel in **Fig. 1A**, corresponding to the purple mask in the middle panel). Meanwhile, we obtained the global BOLD signal by extracting and averaging BOLD signals from the whole-brain gray matter regions (**Fig. 1A**, the green mask). As shown in a representative subject (**Fig. 1B**), large modulations in the global BOLD signal are often accompanied by corresponding large changes in the CSF signal, suggesting a coupling between the two. To better quantify this BOLD-CSF coupling, we calculated the cross-correlation function between the two, which shows their Pearson’s correlations with different time lags. The resulting BOLD-CSF cross-correlation function is characterized by a positive peak (0.17; *p* < 0.0001, permutation test, see **Methods** for details) at the lag of −6 seconds (i.e., shifting CSF ahead of time by 6 seconds) and a negative peak *(r* = −0.17; *p* < 0.0001, permutation test) at the lag of +3 seconds (**Fig. 1C**, upper panel); intriguingly, this temporal pattern closely resembles the one obtained in the previous study^39^ despite with the lower temporal resolution. The cross-correlation function was also calculated between the negative derivative of the global BOLD signal and the CSF signal. It showed a strong positive peak around the lag of −3 second (0.20; *p* < 0.0001, permutation test), also consistent with the previous finding^39^. Taken together, these results confirmed the existence of a significant BOLD-CSF coupling with specific and reproducible temporal patterns in the ADNI dataset.

**Fig. 1.**
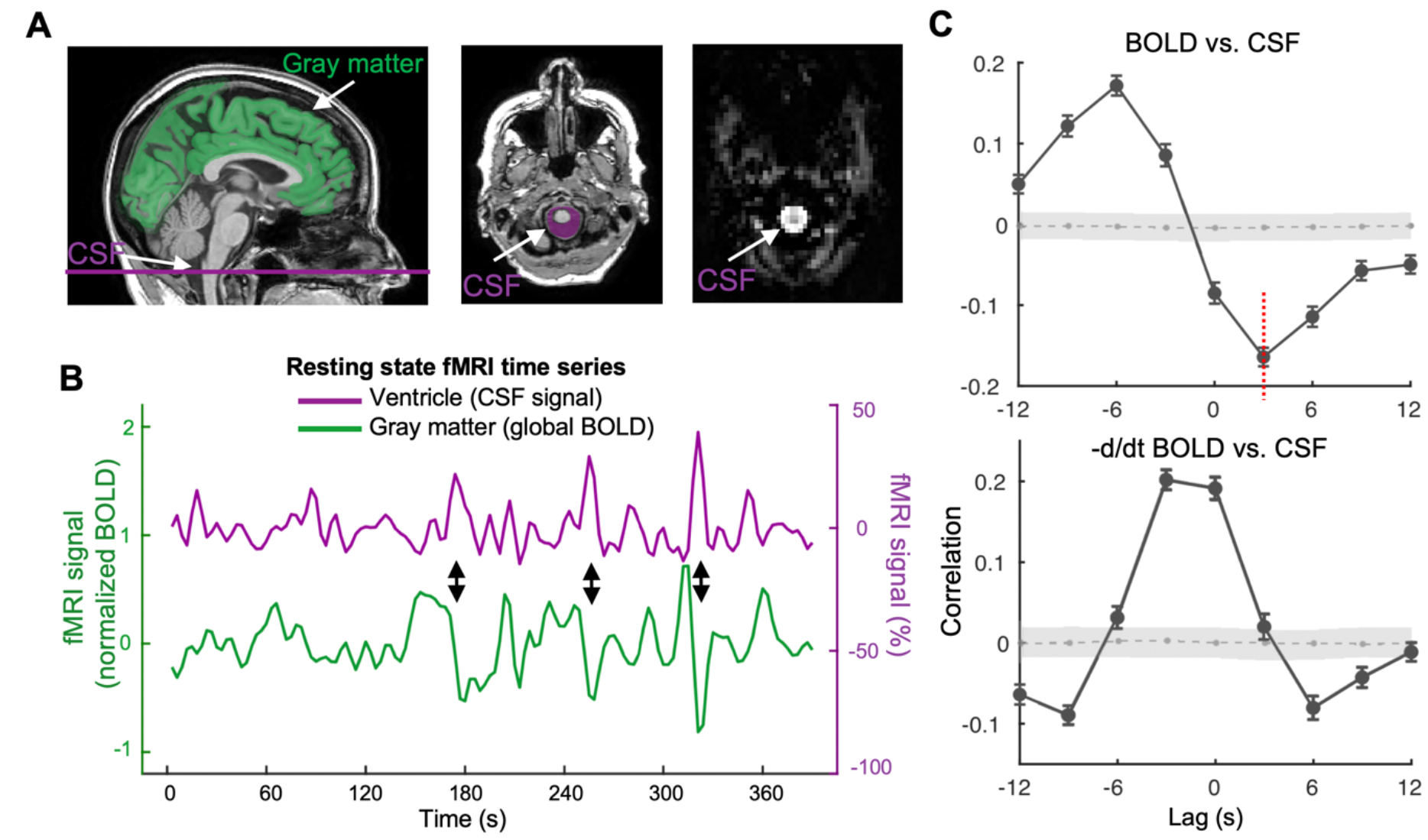
The global BOLD signal is coupled with CSF changes. (**A**) The global BOLD signal was averaged across the gray matter regions (the green mask on an exemplary T1-weighted image in the left panel) whereas the CSF signal was extracted from the CSF regions at the bottom slice of the fMRI acquisition (the middle and right panel). The CSF appears much brighter than the surrounding areas in the T2*-weighted fMRI image (the right panel). (**B**) The global BOLD signal and the CSF signal from a representative subject showed corresponding changes (indicated by black arrows). (**C**) The cross-correlation function between the global BOLD signal and the CSF signal averaged across 158 sessions (upper), and the one between the negative derivative of global BOLD signal and the CSF signal (lower). The gray shaded region denotes 95% confidence intervals calculated with shuffled signals (see **Methods** for details). Gray dashed-line within the shaded region shows mean correlation of the null distribution from permutation test at each time lag. Error bar represents the standard error of the mean (SEM). These cross-correlation functions show a very similar shape to those reported in the previous study. The cross-correlation (−0.17, *p* < 0.0001, permutation test) at the +3 seconds lag (red dashed line), which also showed the strongest coupling in the previous study, was used for quantifying the BOLD-CSF coupling for subsequent analyses.

### Relationships between the BOLD-CSF coupling and AD-related pathology

We then investigated whether the BOLD-CSF coupling is related to the AD-related pathology, including AD risk factors, disease condition, and neurobiological and neuropsychological markers. We used the negative peak BOLD-CSF correlation at the 3-sec lag (**Fig. 1C**, the red dashed line in the upper panel) to represent the strength of the BOLD-CSF coupling, since the previous study with superior imaging quality found the strongest BOLD-CSF correlation at the negative peak around the same lag^39^. The BOLD-CSF coupling strength was first related to the age and gender, two major risk factors for AD. It is significantly correlated with the age (Spearman’s *r* = 0.24; *p* = 0.011 based on a linear mixed effect model with Satterthwaite’s method, the same hereinafter unless noted otherwise; see **Methods** for details) with older subjects showing relatively weaker (less negative) BOLD-CSF coupling (**Fig. 2A**). The BOLD-CSF coupling also appeared to be significantly stronger *(p* = 0.026) in males than in females (**Fig. 2B**). We then compared the BOLD-CSF coupling in subjects under different disease conditions after adjusting for age and gender. The BOLD-CSF coupling strength exhibited a dose-response relationship with the disease condition, lower *(p* = 0.035) from the HC > SMC > MCI > AD groups (**Fig. 2C**). The age- and gender-adjusted BOLD-CSF coupling was also compared across different apolipoprotein E (APOE) gene carriers and showed a marginally significant *(p* = 0.077) dependency on the copies of APOE ε4 alleles (**Fig. 2D**). The results regarding the disease condition and APOE gene could be significantly underpowered due to limited AD patients (N = 7) and 2 APOE ε4 alleles carriers (N = 11). We thus augmented the sample by adding 22 AD and 14 HC subjects, 5 of whom carry 2 APOE ε4 alleles, from another subset of the ADNI dataset (see **Methods** section for details about data selection). The same analysis on this augmented sample (see the sample characteristics in **Table S1**) led to similar findings with improved statistical significance, particularly for the difference *(p* = 0.0078) between the AD and HC groups (**Fig. S1C**). The relationship between the BOLD-CSF coupling and the APOE ε4 copies also reached the statistical significance *(p* = 0.041) with the augmented sample size (**Fig. S1D**). It is also worth noting that the relationships of the BOLD-CSF coupling to age *(p* = 0.020) and gender *(p* = 0.032) remained significant with controlling for disease condition (**Fig. S2**). In summary, the BOLD-CSF coupling was gradually weaker from the HC to SMC, to MCI, and then to AD groups, conditions known to show a gradually increasing severity of AD-related symptoms; it also appeared to be weaker in older subjects, females, and APOE ε4 gene carriers, who are known to have a higher risk of developing AD pathology.

**Fig. 2.**
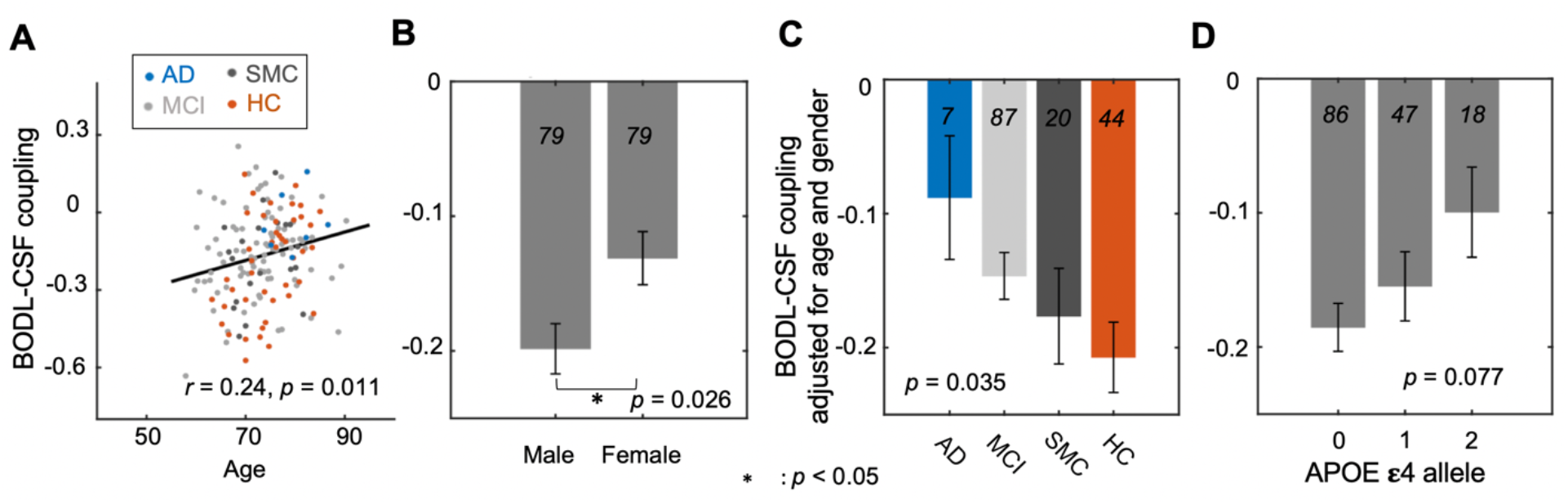
The dependency of the BOLD-CSF coupling on AD risk factors and disease conditions. (A) The strength of the BOLD-CSF coupling, quantified as the correlation between the global BOLD signal and CSF at +3 sec lag, shows a significant correlation (Spearman’s *r* = 0.24, *p* = 0.011, the linear mixed model with Satterthwaite’s method) with age across the 158 sessions. (B) Male participants showed a larger amplitude of the BOLD-CSF coupling as compared with females *(p* = 0.026). (C) The BOLD-CSF coupling, after adjusting for age and gender, decreases gradually *(p* = 0.035) from the healthy controls (HC), to significant memory concern (SMC), to the mild cognitive impairment (MCI), and then to Alzheimer’s Disease (AD) group. (D) The age- and gender-adjusted BOLD-CSF coupling is also marginally *(p* = 0.077) correlated with the APOE ε4 allele. Error bar in this figure represents the SEM. The sample sizes of each sub-groups are shown by numbers on the bars of the bar plots. The analysis was repeated for an augmented sample with more AD patients and healthy controls, and see Fig. S1 for the results.

The BOLD-CSF coupling was then correlated with neurobiological and neuropsychological markers more specifically related to AD pathology, including the cortical Aβ level as measured by the florbetapir standardized uptake value ratio (SVUR) averaged over the entire cortex^40^ and the cognitive performance as measured by the MMSE score. After adjusting the age and gender, a significant positive correlation (Spearman’s *r* = 0.20, *p* = 0.019) was found between the BOLD-CSF coupling and the cortical Aβ level measured at baseline around the same time (**Fig. 3A**). Thus, subjects showing weaker BOLD-CSF coupling exhibited more Aβ accumulated in the cortex. We did not, however, find a significant relationship between the BOLD-CSF coupling and the Aβ accumulation in the subsequent two years (**Fig. 3B**). Rather, the two-year longitudinal change of the MMSE score (**Fig. 3D**), but not its baseline value (**Fig. 3C**), is negatively correlated (Spearman’s *r* = −0.20, *p* = 0.013) with the negative BOLD-CSF coupling, suggesting the association between the weak BOLD-CSF coupling and large cognitive decline over the following two years. These significant correlations did not seem to be driven by the difference between the AD and HC groups in the BOLD-CSF coupling because they remained significant *(p* < 0.05) after removing these two groups from the analysis (**Fig. S4**). We also repeated the above analyses by using another reference region (i.e., eroded white matter) for Aβ SUVR calculation (**Fig. S5**) or across subjects rather than sessions (see **Methods** section for details), and the major results remain unchanged (**Fig. S6**). We examined how the BOLD-CSF correlations at different time lags may be related to the Aβ and cognition decline, and the results showed that they were only significant *(p* < 0.05) at time lags showing large negative BOLD-CSF coupling (i.e., at lag 0-sec and +3-sec; **Fig. S7**). The relationships of the BOLD-CSF coupling with age, gender, disease condition, and APOE gene were re-assessed while controlling for the cortical Aβ level, and the BOLD-CSF coupling remained significantly correlated with age *(p* = 0.049) but not for gender *(p* = 0.06), disease condition *(p* = 0.32), and APOE gene *(p* = 0.86) (**Fig. S3**).

**Fig. 3.**
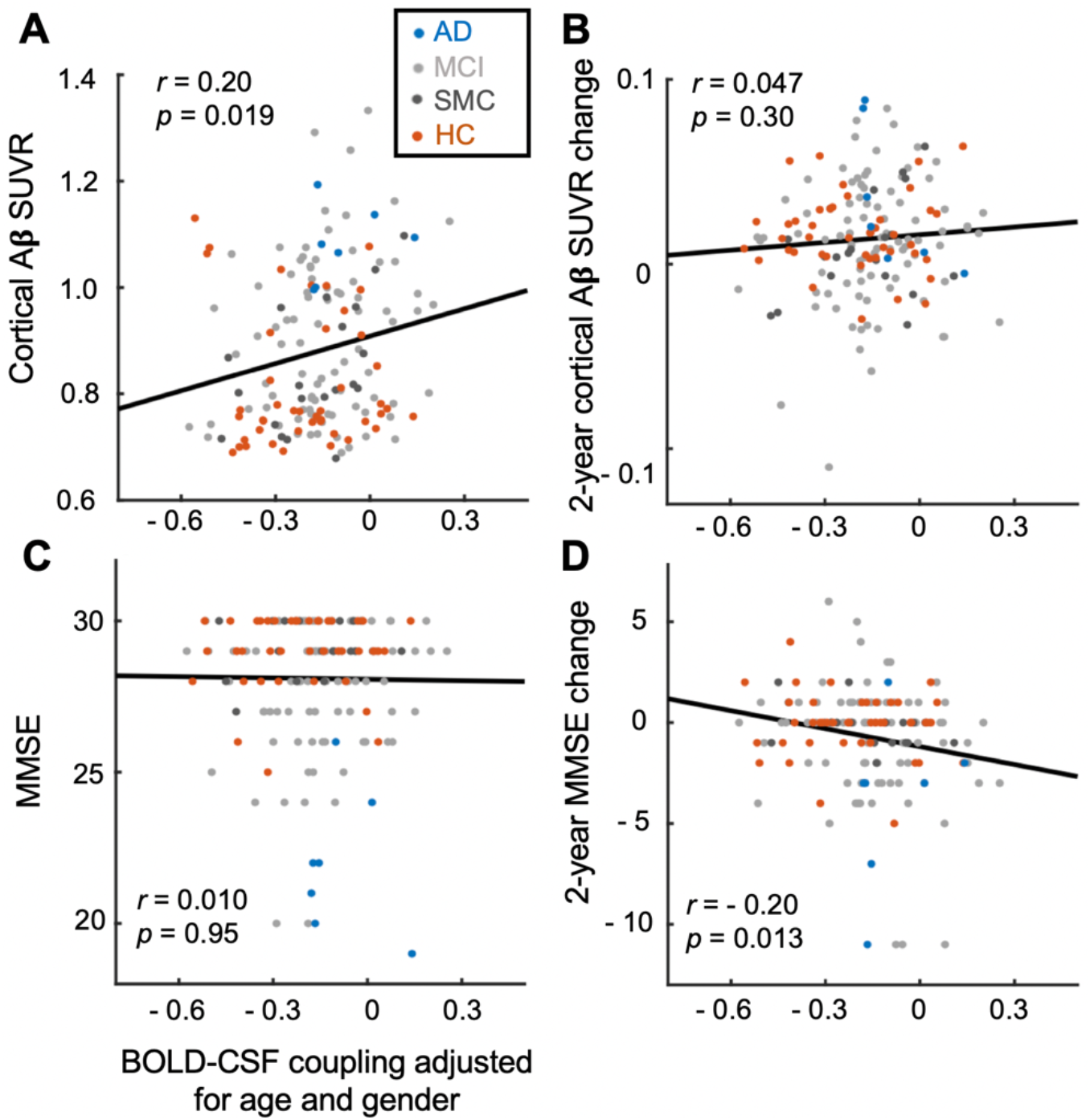
The BOLD-CSF coupling is correlated with the cortical Aβ and cognitive decline. (**A-B**) The BOLD-CSF coupling adjusted for age and gender is significantly correlated (Spearman’s *r* = 0.20, *p* = 0.019, N = 158) with the cortical Aβ SUVRs at baseline (left) but not their changes in the following two years (right). (**C-D**) The BOLD-CSF coupling adjusted for age and gender is significantly correlated (Spearman’s *r* = −0.20, *p* = 0.013, N = 158) the MMSE score changes in the following two years (right) but not with its baseline value (left). The linear regression lines were estimated based on the least-squares fitting^78^. Each dot represents a single session. AD, MCI, SMC, and HC sessions are colored with blue, light gray, dark gray, and orange, respectively.

### The BOLD-CSF coupling’s relationship with AD pathology is not dependent on the global signal amplitude

Next, we sought to understand whether the association between the BOLD-CSF coupling and AD pathology is mediated by the change of global BOLD signal amplitude. Converging evidence has suggested that the fluctuation amplitude of global BOLD signal^20,21,27–30,39^, as well as its coupling with CSF^39^, is highly dependent on sleep/wake state, and much stronger during sleep than wake. Thus, the weak BOLD-CSF coupling may simply reflect a tendency of staying awake inside the scanner for subjects with AD-related pathology. To test this possibility, we investigated how the global BOLD amplitude, i.e., the fluctuation amplitude of the global BOLD signal, may affect the relationship between the BOLD-CSF coupling and AD-related markers. Consistent with the previous study^39^, a stronger BOLD-CSF coupling tends to be associated with *(p* = 0.00062) a larger global BOLD amplitude (**Fig. 4A**) indicative of a drowsy state or sleep^21,30^. However, the global BOLD amplitude is not significantly correlated with either the baseline cortical Aβ *(p* = 0.13) or the longitudinal MMSE changes *(p* = 0.40) (**Fig. 4B** and **4C**). Moreover, after adjusting for the global BOLD amplitude, the relationships of the BOLD-CSF coupling with the baseline Aβ and the longitudinal MMSE changes remained significant *(p* = 0.043 and 0.020 respectively, **Fig. 4D** and **4E**). We also repeated the above analysis by replacing the global BOLD amplitude with an fMRI-based arousal index^41,42^. Although the arousal index estimates the arousal level through distinct information, i.e., the co-activation patterns of individual fMRI volumes, it showed a strong correlation with the global BOLD amplitude (*r* = 0.503, *p* = 1.55×10^-10^, N = 158). Similarly, controlling for the effect of the arousal index did not affect the relationship of the BOLD-CSF coupling with either the baseline Aβ *(p* = 0.022) or the longitudinal MMSE changes *(p* = 0.0094) (**Fig. S8**). Therefore, the weakening of the BOLD-CSF coupling associated with the Aβ accumulation and cognitive decline is unlikely due to a reduced global BOLD amplitude that is suggestive of a brain state change, but reflects a true disruption of the coupling between the global brain signal and CSF flux.

**Fig. 4.**
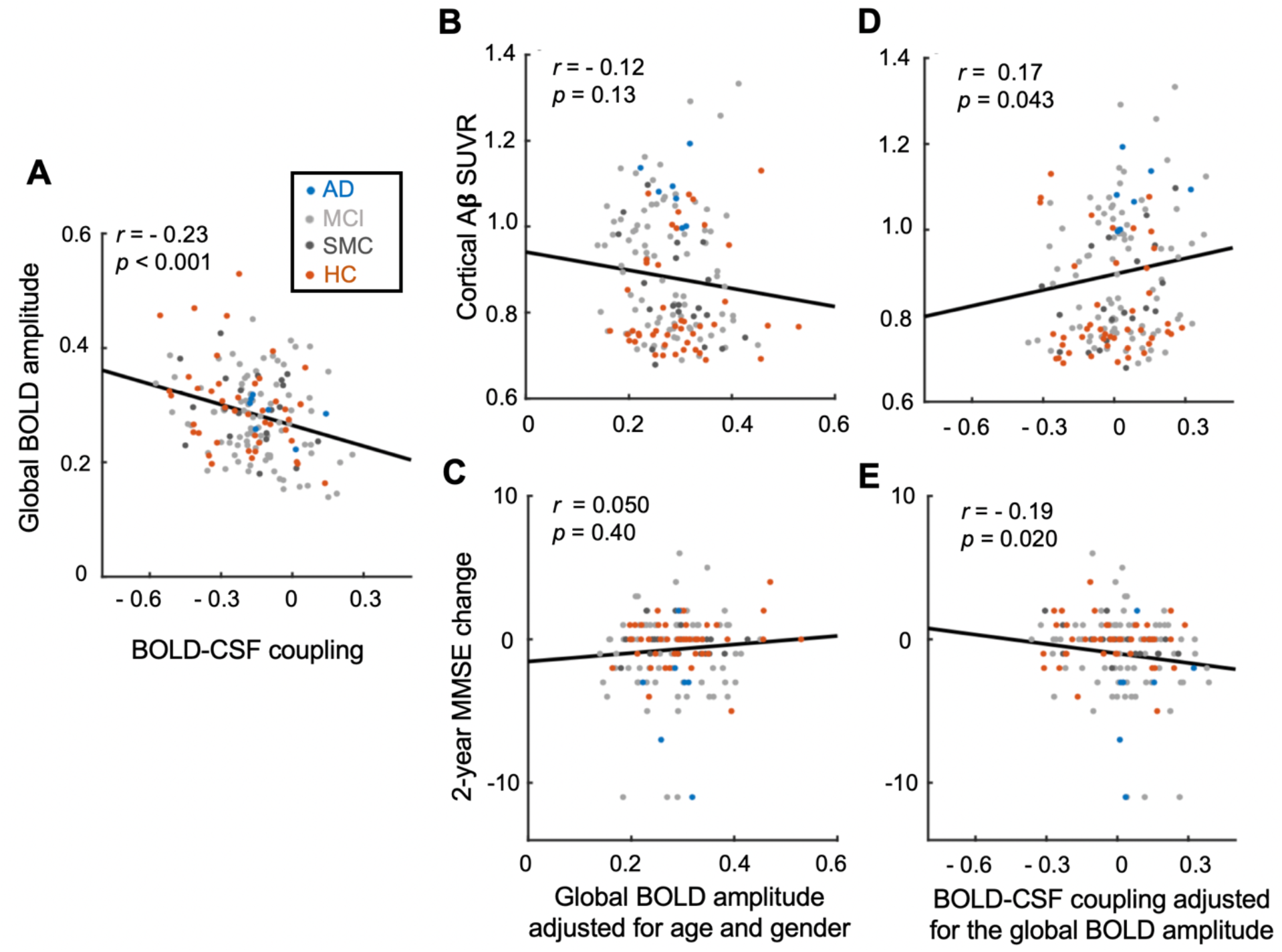
The role of the global BOLD amplitude in the relationships between the BOLD-CSF coupling and AD-related markers. (**A**) The strength of the BOLD-CSF coupling is dependent (Spearman’s *r* = −0.23, *p* < 0.001, N = 158 sessions) on the fluctuation amplitude of the global BOLD signal after adjusting for age and gender. (**B-C**) The amplitude of the global BOLD signal, adjusted for age and gender, is not significantly correlated with either the cortical Aβ level (**B**) or the 2-year longitudinal change of MMSE score (**C**). (**D-E**) The BOLD-CSF coupling remains significantly correlated with the cortical Aβ level (**D**) and the MMSE changes (**E**) after adjusting for age, gender, and global BOLD amplitude. Each dot represents a single session. AD, MCI, SMC, and HC sessions are colored with blue, light gray, dark gray, and orange, respectively.

## Discussion

We show a strong coupling between the global brain BOLD signal and CSF flux in the rsfMRI data from older adults in the ADNI project. The strength of this BOLD-CSF coupling varied considerably across subjects and is significantly weaker in those with a higher risk for AD or having developed AD-related diseases. Importantly, the BOLD-CSF coupling strength is significantly associated with the cortical Aβ level and the MMSE score change in the subsequent two years, indicating its close relationship with the AD pathophysiology. These findings suggest an important role of the neural and physiological modulations associated with the large global brain signal in Aβ clearance and thus the AD pathogenesis, presumably through their involvement in the glymphatic process.

The global brain signal and CSF flux become weakly coupled in subjects with AD-related diseases or at higher risk of developing AD. A gradual weakening of the BOLD-CSF coupling is evident from the HC, to SMC, to MCI, and then to AD groups demonstrating increasing severity of AD-related symptoms. The BOLD-CSF coupling also appeared significantly weaker in older subjects and females, who are known to have a higher risk of developing AD^43,44^, and the differences remain significant with controlling for the disease condition effect. The BOLD-CSF coupling is also weaker in APOE gene carriers, but this association became non-significant after adjusting for disease condition. The dependency of the BOLD-CSF coupling on the disease condition, as well as the APOE gene, can be at least in part attributed to its dependency on cortical Aβ, because they are no longer significant when controlling for Aβ levels. The association between the BOLD-CSF coupling and the longitudinal MMSE change may also be related to the cortical Aβ accumulation, which appears earlier than the cognitive decline in the AD pathogenesis^45,46^. Therefore, the key to understanding the link of BOLD-CSF coupling with the AD-related pathology lies in its relationship with the cortical Aβ level, which is likely related to the Aβ clearance through the glymphatic system.

The BOLD-CSF coupling could be related to the characteristic global brain activity at transient arousal modulations. Despite the lower spatial and temporal resolutions than the previous study^39^, the ADNI data displayed a consistent BOLD-CSF coupling as evidenced by the same pattern of the cross-correlation functions^39^. The BOLD-CSF coupling has been shown to be present during sleep with the global BOLD signal showing large-amplitude fluctuations^39^. Consistent with this finding, we observed stronger BOLD-CSF coupling at larger global signal amplitudes (**Fig. 4A**). The dependency of the global BOLD signal on the brain state, particularly its large fluctuations during sleep, has been repeatedly reported by rsfMRI studies^27,29–35^. However, not until recently did we have more clues about the neurophysiological underpinnings of this phenomenon. A recent multimodal study on monkeys and humans suggested that the large-amplitude global BOLD fluctuation is driven by global brain co-activations that are associated with a characteristic neurophysiological event, i.e., the sequential spectral transition (SST) event^21^. Importantly, both the spatial pattern of the global co-activations, which showed specific deactivations in subcortical arousal-regulating regions^21^, and the time-frequency pattern of the SST, which displayed a transition from wake-dominant alpha-beta (8-30 Hz) activity to sleepdominant delta (0.5-4 Hz) activity^20,21^, indicated a transient modulation in brain arousal. Thus, the large global BOLD signal, as well as its coupling with CSF, could be related to transient arousal modulations in the drowsy state or during sleep.

The arousal-related global brain co-activations are accompanied by physiological modulations that can potentially drive CSF movements with the involvement of the autonomic system. First, the arousal-related global brain activity is likely associated with slow (< 0.1 Hz) but large arterial constrictions/dilations that may drive the CSF fluxes. The arterial pulsation has been regarded as the major driving force that propels CSF influxes along the periarterial spaces^6,14–16^, but it is expected to decrease during sleep due to reduced physical activity and thus cannot explain the greatly enhanced glymphatic process during sleep. Reduced tissue resistance, as evidenced by decreased interstitial spaces, was thus proposed to explain the sleep-related enhancement of glymphatic process^4^ that maybe necessary but not sufficient for sleep-related CSF flux. The vascular modulations at the arousal-related global brain activity may further facilitate this process by promoting large-scale CSF movements. The global BOLD signal of brain tissues is correlated with peripheral cardiac signals (envelop amplitude and heart rate variability)^47–50^ and also preceded by strong, anti-phase BOLD changes in the internal carotid arteries (ICA)^51^. Given the saturated oxygenation in arteries, the ICA signal changes are likely attributed to cerebral blood volume changes caused by arterial constrictions/dilations. The cardiac-related global BOLD component, which is related to transient arousal changes during sleep^52,53^, also showed a systematic delay from the gray matter ventromedially towards deep white matter structures^36^. Such a delay was well explained by a model assuming the modulation of pial arteries by sympathetic innervation from the superior cervical ganglion^36^. Thus, the autonomic sympathetic modulation at transient arousal modulations may serve as the driving forces of arterial tone modulations, which in turn propel the CSF movements. Second, the arousal-related global brain activity is also accompanied by large respiratory modulations^38,54^, which have been hypothesized to be the second driving force of CSF movements^17,18^. Therefore, the autonomic modulation at the transient arousal fluctuation during sleep might also drive the CSF flux through its effect on respiration. However, it is also possible that the respiratory change is simply another correlated event at transient arousal modulations.

The neural and physiological changes at transient arousal modulations may further facilitate the glymphatic clearance in two hypothetical ways. First, they might facilitate CSF movements by affecting its production. The superior cervical ganglia are known to have strong sympathetic inhibitory control over the choroid plexuses in the brain ventricles^55^, which are the major brain structure to produce CSF. The autonomic modulations associated with the global brain coactivations are thus expected to also affect the CSF production that may in turn affect its movement. Second, the coordinated CSF movement and global brain co-activations might be another factor facilitating brain waste clearance. The arousal-related global brain activity showed a much larger amplitude in the motor/sensory regions^21^, where the Aβ accumulates much slower than the other higher-order cortical regions in the AD prognosis^56^, suggesting a potential interaction effect between the brain activation and the glymphatic process. However, these two potential mechanisms are purely conjectural. They remain to be investigated by future studies, particularly those on animal models.

The co-modulation of the BOLD-CSF coupling and the cortical Aβ should not be attributed to the change of global BOLD signal amplitude suggestive of the brain state change, nor interpreted as a causal relationship. The BOLD-CSF coupling is significantly correlated with the global BOLD signal amplitude, as well as the fMRI-based arousal index, confirming its dependency on the brain state. However, the global signal amplitude itself is not correlated with the cortical Aβ level, nor did the control of this factor affect the association between the BOLD-CSF coupling and Aβ. Thus, the reduced BOLD-CSF coupling at the high Aβ level likely reflects a disrupted coupling between the global brain activity and CSF movements. At the same time, this correlation alone cannot tell us whether the disrupted BOLD-CSF coupling is a cause or a consequence of the Aβ accumulation. It is worth noting that the BOLD-CSF coupling is correlated with age, even controlling for cortical Aβ level. However, this piece of evidence is insufficient to exclude the possibility that the increased cortical Aβ, which is often deposited in the perivascular spaces to result in cerebral amyloid angiopathy^57,58^, may block CSF flux coupled with the global brain activity and thus disrupt the BOLD-CSF coupling. It is also possible that the decrease of BOLD-CSF coupling and the accumulation of cortical Aβ are two processes facilitating each other after either one initiating their interaction. Testing these possibilities remains a challenge for future work.

In summary, we found a strong coupling between the global BOLD signal and the CSF inflow in the ADNI rsfMRI data. More importantly, the strength of this BOLD-CSF coupling was significantly associated with various factors related to AD pathology, including the cortical Aβ level. These findings provide initial evidence for the role of large-scale neural and physiological modulations, which are associated with transient arousal modulations in the drowsy state or during sleep, in the glymphatic clearance of toxic brain wastes, including Aβ. They also suggest that the coupling of the global brain signal and CSF inflow may serve as a potential imaging marker for clinical evaluation.

## Methods

### Participants and Study data

This is an analytic observational study to investigate the potential relationship between the coupling between the global brain activity and CSF inflow and the AD-related pathology. We included 118 subjects from the ADNI project (ADNI-GO, ADNI-2, and ADNI-3), based on the availability of rsfMRI, longitudinal MMSE scores, and longitudinal 18F-AV45 amyloid-PET. Specifically, we selected the subjects who had rsfMRI at baseline and amyloid PET and MMSE data at baseline and in a 2-year follow-up. The group include AD (N = 7), mild cognitive impairment (MCI, early MCI and late MCI included; N = 62), significant memory concern (SMC; N = 18), and healthy control (HC; N = 31) subjects as defined by ADNI (http://adni.loni.usc.edu/study-design/), and no subjects in the present study experienced changes in the disease condition over the 2-year period. Among them, 113 subjects also have available APOE genotype information. **Table 1** summarized the subject characteristics, including age at baseline, gender, and numbers of APOE ε4 allele carrying as well as corresponding longitudinal MMSE scores and PET-SUVRs. Since the sample size determined by the above inclusion criteria was too small for the AD group, we also augmented the data by another subset of ADNI data used for a different research project. This second subset of ADNI data includes subjects with longitudinal rsfMRI data at two time points separated by two years, but not all of the had the corresponding longitudinal amyloid-PET or MMSE data. 23 AD and 18 HC sessions from 22 AD and 14 HC subjects were added into the original sample to create an augmented sample (see **Table S1** for the subject characteristics of the augmented sample), which was only used for reexamining the effects of age, gender, disease condition, and APOE gene on the BOLD-CSF coupling. All subjects provided written informed consent, and investigators at each ADNI participating sites obtained ethical approval from the corresponding Institutional Review Board (IRB).

The rsfMRI, MMSE, APOE genotype, and amyloid-PET data at baseline were obtained in the same study visit (the “visit” was defined by ADNI, see “visit code” details in https://adni.loni.usc.edu/wp-content/uploads/2008/07/inst_about_data.pdf). The MMSE and amyloid-PET data acquired in a 2-year follow-up (25.0 ± 1.5 months interval for the MMSE interval, 23.9 ± 1.2 months interval for the amyloid-PET) were also used. Amyloid-PET SVUR data, MMSE scores, and APOE gene genotypes were directly obtained from “UC Berkeley – AV45 Analysis [ADNI1,GO,2,3] (version: 07_28_19)” ^59,60^, “Mini-Mental State Examination (MMSE) [ADNI1,GO,2,3]” ^61^, and “APOE – Results [ADNI1,GO,2,3]” respectively from the ADNI website (http://adni.loni.usc.edu/). There are a total of 158 sessions from the 118 subjects having the baseline rsfMRI and 2-year longitudinal MMSE and amyloid-PET data. Twenty-nine subjects have two rsfMRI sessions at years 0 and 2, as well as the MMSE and amyloid-PET at years 0, 2, and 4; four subjects with 3 rsfMRI sessions (for each subject) with corresponding 4 sessions of MMSE and amyloid-PET (two-year interval); and one subject with 4 fMRI sessions and 5 corresponding MMSE and amyloid-PET sessions. Overall, our data included 7 sessions from 7 AD patients, 87 sessions from 62 MCI subjects, 20 sessions from 18 SMC subjects, and 44 sessions from 31 HC subjects (see **Table 1**). The APOE genotype data are available for 151 sessions from 113 subjects.

The use of de-identified data from the ADNI and the sharing of analysis results have been reviewed by the Pennsylvania State University IRB and also strictly followed the ADNI data use agreements.

### Image acquisition and preprocessing

All rsfMRI data were acquired by 3 Tesla MR scanners in different ADNI participating sites following the same protocol (http://adni.loni.usc.edu/methods/documents/mri-protocols/). Each rsfMRI session began with an MPRAGE sequence (echo time (TE) was minimum full echo time, repetition time (TR) = 2300ms), which was used for anatomical segmentation and template normalization^62^ (see acquisition details in http://adni.loni.usc.edu/methods/documents/). For the rsfMRI, 140 (ADNI-GO and ADNI-2) or 200 (ADNI-2 (Extended fMRI sessions) and ADNI-3) fMRI volumes were obtained with a 3D echo-planar image (EPI) sequence (ADNI-GO&ADNI-2: flip angle = 80°, spatial resolution = 3×3×3 mm^3^, slice thickness = 3.3 mm; ADNI-3: flip angle = 90°, slice thickness = 3.4 mm; see http://adni.loni.usc.edu/methods/documents/ for details) with TR/TE = 3000/30ms. Amyloid PET data were acquired from the 50 ~ 70 minutes post-injection of florbetapir-fluorine-18 (^18^F-AV-45) (see the detailed description in http://adni.loni.usc.edu/data-samples/data-types/).

For the rsfMRI data, we applied the preprocessing procedures typical for rsfMRI signals using a 1000 Functional Connectomes Project script (version 1.1-beta; publicly available at https://www.nitrc.org/frs/?group_id=296) with some minor modifications (Biswal et al. 2010). Specifically, we performed motion correction, skull stripping, spatial smoothing (full width at half maximum (FWHM) = 4mm), temporal filtering (bandpass filter, 0.01~ 0.1 Hz), and linear and quadratic temporal trends removal. The first 5 and last 5 rsfMRI volumes were discarded to allow for the magnetization to reach steady state and also to avoid the edge effect of the temporal filtering. We then co-registered each fMRI session to corresponding high-resolution anatomical images (T1-weighted MRI) and then to the 152-brain Montreal Neurological Institute (MNI-152) space. Our modifications of the original scripts include performing spatial registration between functional and anatomical images using align_epi_anat.py ^63^ in AFNI software and resampling the preprocessed fMRI data into a spatial resolution of 3×3×3 mm^3^. We also skipped the nuisance regression of global signal, CSF signal, motion parameters to keep consistent with the previous study of the BOLD-CSF coupling^39^. The global brain signal and the CSF signal are the major interests of our study. The head motion, which had been suggested to cause rsfMRI changes^64^, was recently found to be more likely a by-product associated with, rather than the cause of, the arousal-related global BOLD signal^38^. We thus skipping the motion parameter regression to avoid attenuating the global BOLD signal.

For the Amyloid PET data, we directly used SUVR data summarized in “UC Berkeley – AV45 Analysis [ADNI1,GO,2,3] (version: 07_28_19)”^59^ (available at https://ida.loni.usc.edu/). Briefly, the florbetapir images were averaged, aligned spatially, and interpolated to a standard voxel size. The images were then spatially smoothed and then reregistered to the space of T1-weighted MRI in attempted to extract mean florbetapir uptake at gray matter. Notably, we used the composite region, including the whole cerebellum, brainstem/pons, and subcortical WM regions, as the reference region^40^, and the cortical amyloid-β (Aβ) SUVR was calculated as the ratio of the mean florbetapir uptake at the gray matter and that at the composite reference region.

### Extraction of the global BOLD signal and the CSF signal

We extracted the global BOLD signal from the gray matter regions of the cortex. A mask of the gray matter regions was defined based on the Harvard-Oxford cortical and subcortical structural atlases^65^ (https://neurovault.org/collections/262/). We transformed the gray-matter mask from the MNI-152 space back to the original space of each subject/session to avoid spatial blurring from the registration process with the same rationale to a previous study^39^. For each session, the fMRI signals were extracted from and averaged within the individual’s gray-matter mask after all the pre-processing procedures but before the spatial registration to the MNI-152 space. The fMRI signals were also normalized to Z-score at each voxel before being averaged. Therefore, all the voxels have the equal fluctuation amplitude, and the amplitude of their average, i.e., the global BOLD signal, would reflect the level of global synchronization.

The CSF signals were extracted from the bottom slices of fMRI acquisition, which are near the bottom of the cerebellum for all the subjects, to obtain high sensitivity to the CSF inflow effect as described previously^39^. For each session, a CSF mask was manually defined at the bottom slice based on the T2*-weighted fMRI image, and its location was further confirmed with the T1-weighted structural image. The fMRI signals were extracted from and averaged within the individual’s CSF mask after several pre-processing procedures, including motion correction, skull stripping, and temporal filtering, but before the spatial registration to the MNI152 space.

### The coupling between the global BOLD signal and the CSF signal

We calculated the cross-correlation functions between the global BOLD signal and the CSF signal obtained through the above procedures to quantify their coupling, similarly to the previous study^39^. The cross-correlation functions show the Pearson’s correlations between the global BOLD signal and the CSF signal at different time lags. While the negative peak at the lag of +3 second is of a similar amplitude as the positive peak at the lag of −6 second in our study, it appeared to be stronger than the positive peak in the previous study with higher spatial and temporal correlations^39^. We thus used the BOLD-CSF correlation at this negative peak (the lag of +3 second) for quantifying the strength of the BOLD-CSF coupling. Following the previous study^39^, we also calculated the cross-correlation function between the negative (temporal) derivative of the global BOLD signal and the CSF signal. To test the statistical significance of the BOLD-CSF correlations, we used a permutation method to create a null distribution for the BOLD-CSF correlation. Specifically, we re-calculated cross-correlation functions between the mismatched global BOLD signal and CSF signal after randomly permuting the session ID of the global BOLD signals. This process was repeated by 10,000 times to build the null distribution for the mean BOLD-CSF cross-correlation functions. The p-value for the BOLD-CSF correlations at different lags was obtained by comparing them against the null distribution.

### The relationship between the BOLD-CSF coupling and AD-related pathology

We quantified the BOLD-CSF coupling strength using their cross-correlation at the lag of +3 seconds (the negative peak of the cross-correlation function). We then related the BOLD-CSF coupling to age, gender, disease condition, and APOE gene. The association between the BOLD-CSF coupling and age was investigated across sessions (N = 158), and it was also compared across different sub-groups, e.g., males versus females, using a linear mixed effect model. For the disease condition, we also tested whether the BOLD-CSF coupling strength (age- and gender-adjusted) changed with the severity of the AD-related symptoms (see details about the severity and staging of four conditions at http://adni.loni.usc.edu/study-design/) (**Fig. 2C**). For APOE gene, we also tested the relationship between the BOLD-CSF coupling strength (age- and gender-adjusted) and the number of APOE ε4 alleles. We repeated the same analysis for the augmented dataset with more AD patients and APOE ε4 alleles carriers (**Fig. S1**).

To investigate whether the relationships of the BOLD-CSF coupling with age, gender, and APOE genotypes are driven by the differences between different disease groups, i.e., AD, MCI, SMC, and HC groups, we repeated the above analyses with controlling for the disease condition (**Fig. S2**). To test whether the relationship between the BOLD-CSF coupling and age/gender/APOE ε4 allele number/disease condition was related to the cortical Aβ SUVR level, we repeated the above analyses by controlling for the cortical Aβ SUVR at baseline (**Fig. S3**).

We also examined the associations between the BOLD-CSF coupling and AD-related markers, including the cortical Aβ level ^66–68^ and MMSE scores ^69–71^ at the baseline and their changes during the subsequent 2 years. The BOLD-CSF coupling strength was adjusted for age and gender effect before all subsequent analyses. The Aβ deposition at the gray matter regions was estimated with the whole-cortical Aβ SUVR, and we also estimated its 2-year change by subtracting its baseline value from that of the 2-year follow-up. The cortical Aβ SUVR was calculated with respect to two different reference regions, but its association with the BOLD-CSF coupling remained similar (**Fig. 3** and **Fig. S5**). We also correlated the baseline MMSE score and its 2-year longitudinal change with the BOLD-CSF coupling. The above correlational analyses were also conducted across the subjects by discarding some data (**Fig. S6**). Specially, for the subjects with data at more than two time points, we discarded those acquired at year 4 and later.

To exclude the possibility that the reported results are driven by the difference between the AD and HC groups in the BOLD-CSF coupling, we repeated the above correlational analyses by removing sessions from the AD and HC groups (**Fig. S4**). Moreover, we also repeated the above analyses by using the BOLD-CSF correlations at different time lags to represent the BOLD-CSF coupling strength (**Fig. S7**).

### The role of the arousal state in the coupling-marker relationships

Since the arousal fluctuation is a major confounding factor in rsfMRI^21,72,73^ and affects the BOLD-CSF coupling to a large extent ^39^, it would be appropriate to investigate the potential arousal effect on the relationships between the BOLD-CSF coupling and AD-related marker. We calculated the standard deviation (SD) of the global BOLD signal for each session to quantify its fluctuation amplitude, which has been linked to the arousal state with a larger value suggesting a drowsy and sleepy state^31,39^. The global BOLD signal amplitude was adjusted for age and gender and then correlated with the BOLD-CSF coupling strength, the baseline amyloid-PET, and the 2-year MMSE changes. Then, the associations between the BOLD-CSF coupling and the AD-related markers were re-evaluated with further controlling for this global BOLD amplitude (see results in **Fig. 4 D** and **E**).

We also repeated the above analysis (in **Fig. 4**) using another fMRI-based arousal index^74^ that is based on a spatial arousal template and thus utilizes the information independent from the global signal amplitude to estimate the arousal level. Specifically, we calculated the spatial correlation between single rsfMRI volumes and an arousal template^74^ and then took the standard deviation of this time-course to quantify the arousal level of each rsfMRI sessions^38^.

### Statistical analysis

Group comparisons of our sample characteristics (**Table 1** and **Table S1**) were performed using two-sample t-test for continuous measures (i.e., age, MMSE and cortical Aβ SUVR) and Chisquared test for categorical measures (i.e., gender and APOE ε4 allele negative/positive) across groups (AD, MCI, SMC, and HC). Significant differences *(p* < 0.05) for each comparison were marked with a bold format. The significance of the BOLD-CSF correlations was tested with a permutation method that shuffled the correspondence between the global BOLD signal and the CSF signal to build the null distributions for the BOLD-CSF correlation at different time lags, similar to the previous study^39^. Spearman’s correlation was used for quantifying the inter-session and inter-subject associations between different quantities, some of which, e.g., the MMSE scores, showed a non-Gaussian distribution^75^. Because some sessions were collected from the same subject, we employed a linear mixed effect model to assess the statistical significance of various relationships in all session-based analyses. Specifically, subject-specific random intercepts were included to account for the dependency of intra-subject sessions. The linear mixed model was implemented by the “lme4” package of R^76^. The corresponding p-values were computed using the R package “lmerTest” based on the Satterthwaite’s degrees of freedom method ^77^. We also validated the major findings in a subject-based analysis based on a simple linear regression model (**Fig. S6**). In this study, a p-value less than 0.05 was regarded as statistical significance.

## Acknowledgments

We thank the ADNI for data collection and sharing through generous contributions from the following: AbbVie, Alzheimer’s Association; Alzheimer’s Drug Discovery Foundation; Araclon Biotech; BioClinica, Inc.; Biogen; Bristol-Myers Squibb Company; CereSpir, Inc.; Cogstate; Eisai Inc.; Elan Pharmaceuticals, Inc.; Eli Lilly and Company; EuroImmun; F. Hoffmann-La Roche Ltd and its affiliated company Genentech, Inc.; Fujirebio; GE Healthcare; IXICO Ltd.; Janssen Alzheimer Immunotherapy Research & Development, LLC.; Johnson & Johnson Pharmaceutical Research & Development LLC.; Lumosity; Lundbeck; Merck & Co., Inc.; Meso Scale Diagnostics, LLC.; NeuroRx Research; Neurotrack Technologies; Novartis Pharmaceuticals Corporation; Pfizer Inc.; Piramal Imaging; Servier; Takeda Pharmaceutical Company; and Transition Therapeutics. The Canadian Institutes of Health Research is providing funds to support ADNI clinical sites in Canada. Private sector contributions are facilitated by the Foundation for the National Institutes of Health (www.fnih.org). The grantee organization is the Northern California Institute for Research and Education, and the study is coordinated by the Alzheimer’s Therapeutic Research Institute at the University of Southern California. ADNI data are disseminated by the Laboratory for Neuro Imaging at the University of Southern California.

## Author contributions

**X.L&F.H** contributed to the conception, design of the work, and investigation; **F.H, A.B.R&J.C** acquired and processed the data; **F.H, A.B.R, Y.G, L.L, & X.L** contributed to data analysis and visualization; **X.L** devoted the efforts to the supervision, project administration and funding acquisition; **F.H, L.L, O.M.B&X.L.** contributed to drafting the paper; and all authors contributed to editing and reviewing of the paper.

## Data and materials availability

The multimodal data, including subject characteristics, rsfMRI, Amyloid-PET SUVR, APOE genotypes, and MMSE scores, are all publicly available at the ADNI website (http://adni.loni.usc.edu/). The ADNI was launched in 2003 as a public-private partnership, led by Principal Investigator Michael W. Weiner, MD. The primary goal of ADNI has been to test whether serial magnetic resonance imaging (MRI), positron emission tomography (PET), other biological markers, and clinical and neuropsychological assessment can be combined to measure the progression of mild cognitive impairment (MCI) and early Alzheimer’s disease (AD). For up-to-date information, see www.adni-info.org. All the code used in the present study are available from the corresponding author upon request.

## Supplementary Materials

### Supplementary figures

**Fig. S1.**
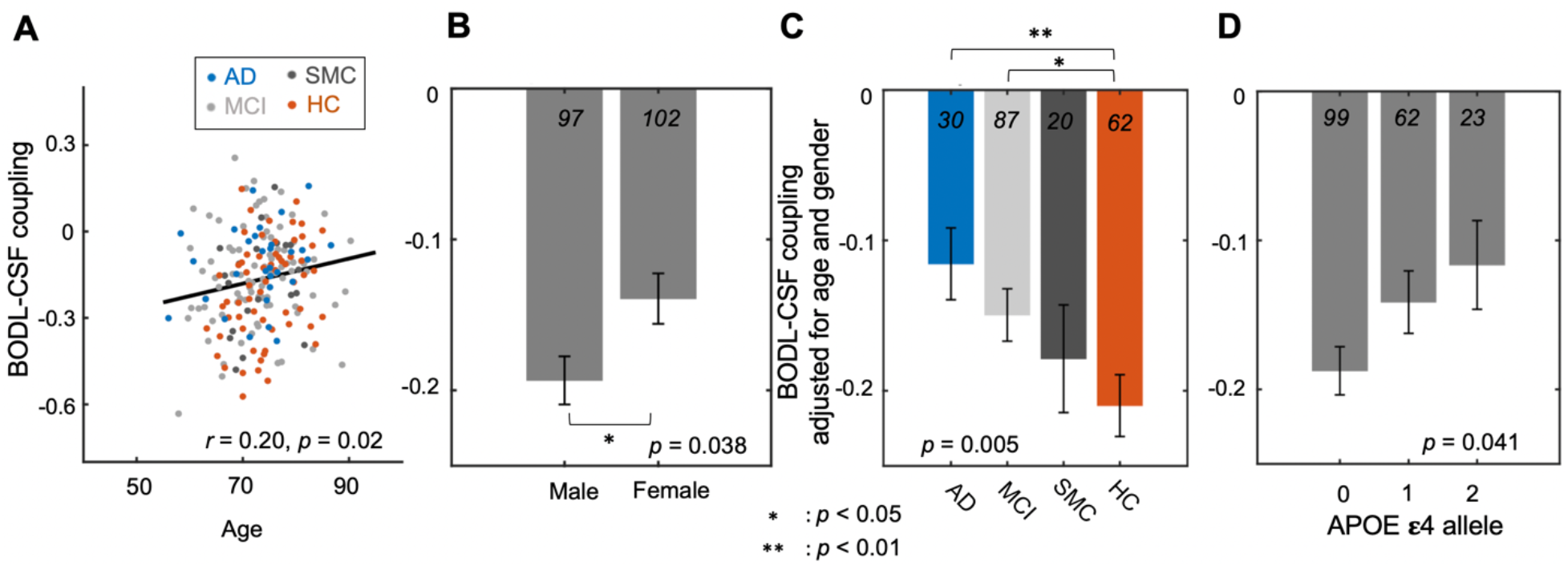
The dependency of the BOLD-CSF coupling on AD risk factors and disease conditions on the augmented dataset. (**A**) When we increased the sample size of AD and HC session (N = 199), the BOLD-CSF coupling shows a significant correlation with age across sessions (Spearman’s *r* = 0.20, *p* = 0.02). AD, MCI, SMC, and HC sessions are colored with blue, light gray, dark gray, and orange, respectively. Each dot represents a session. (**B**) Male subjects have larger amplitudes of this BOLD-CSF coupling compared with female ones *(p* = 0.038) in this augmented dataset. (**C**) The strength of the BOLD-CSF coupling, after adjusting the age and gender effects, also decreases gradually *(p* = 0.005) along with the increased severity of disease condition, i.e., the axis of HC-SMC-MCI-AD. Importantly, significant differences can be found not only between the HC and MCI groups *(p* = 0.045) but also between the HC and AD sessions *(p* = 0.0078). (**D**) The age- and gender-adjusted BOLD-CSF coupling is also significantly correlated with the APOE ε4 allele number (N = 184, *p* = 0.041) across augmented sessions. Error bar in this figure represents the SEM.

**Fig. S2.**
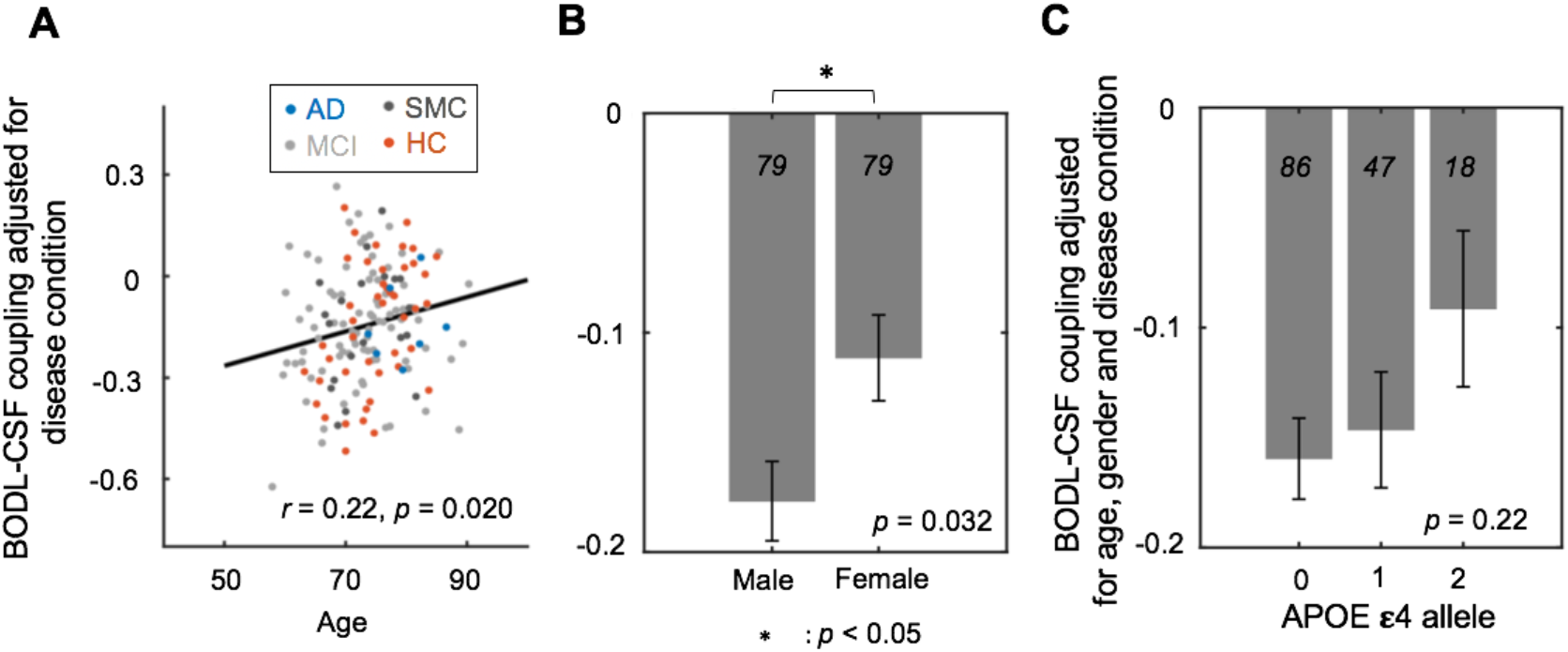
The dependency of the BOLD-CSF coupling on AD risk factors with controlling for disease condition. (**A**) The strength of the BOLD-CSF coupling adjusted for the disease condition (i.e., the effects of AD, MCI, SMC, and CN groups) shows a significant correlation (Spearman’s *r* = 0.22, *p* = 0.020) with age across the 158 sessions. (**B**) Male participants showed a larger amplitude of the BOLD-CSF coupling as compared with females *(p* = 0.032) after controlling for disease condition. (**C**) The age-, gender-, and disease condition-adjusted BOLD-CSF coupling amplitude gradually decreased as the APOE ε4 allele number increases, but this change is not statistically significant *(p* = 0.22). Error bar in this figure represents the SEM.

**Fig. S3.**
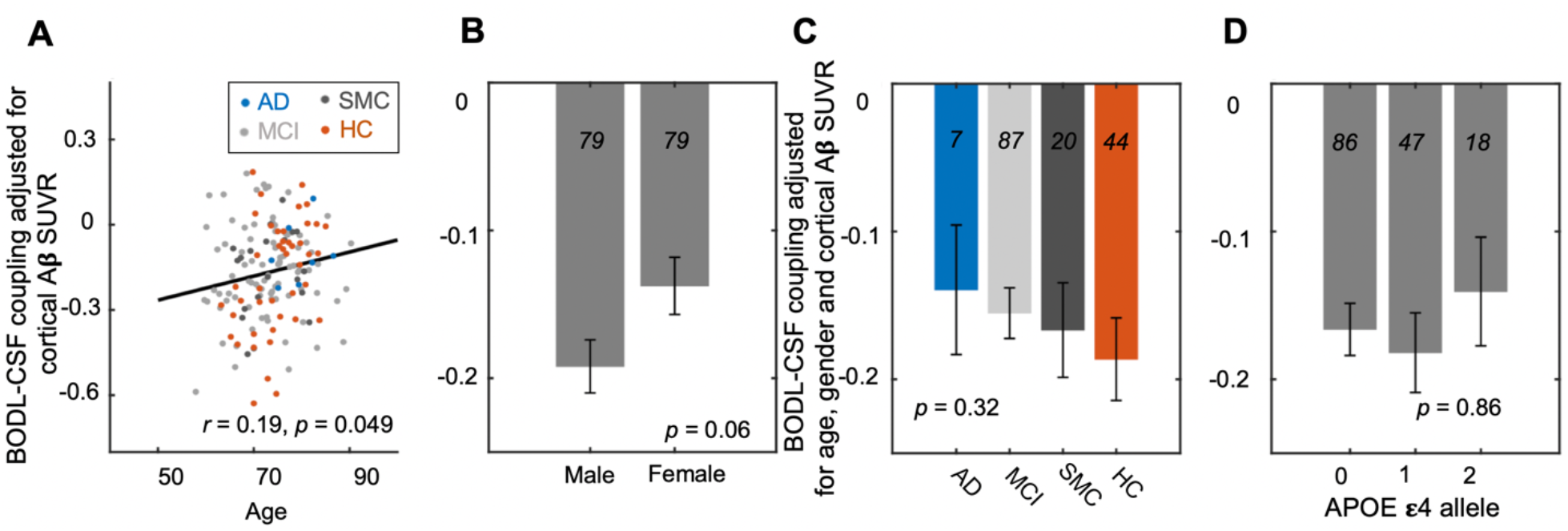
The dependency of the BOLD-CSF coupling on AD risk factors and disease conditions with controlling for cortical Aβ SUVR at baseline. (**A**) The BOLD-CSF coupling adjusted for the cortical Aβ SUVR shows a significant correlation (Spearman’s *r* = 0.19, *p* = 0.049) with age across the 158 sessions. (**B**) Male participants showed a stronger BOLD-CSF coupling (adjusted for cortical Aβ SUVR) as compared with females *(p* = 0.06). (**C**) The BOLD-CSF coupling, after adjusting age, gender, and cortical Aβ SUVR, gradually decrease from the HC, to SMC, to MCI, and then to AD group, but the change is not statistically significant *(p* = 0.32). (**D**) The age-, gender-, and cortical Aβ SUVR adjusted BOLD-CSF coupling is not significantly *(p* = 0.86) correlated with the APOE ε4 allele. Error bar in this figure represents the SEM.

**Fig. S4.**
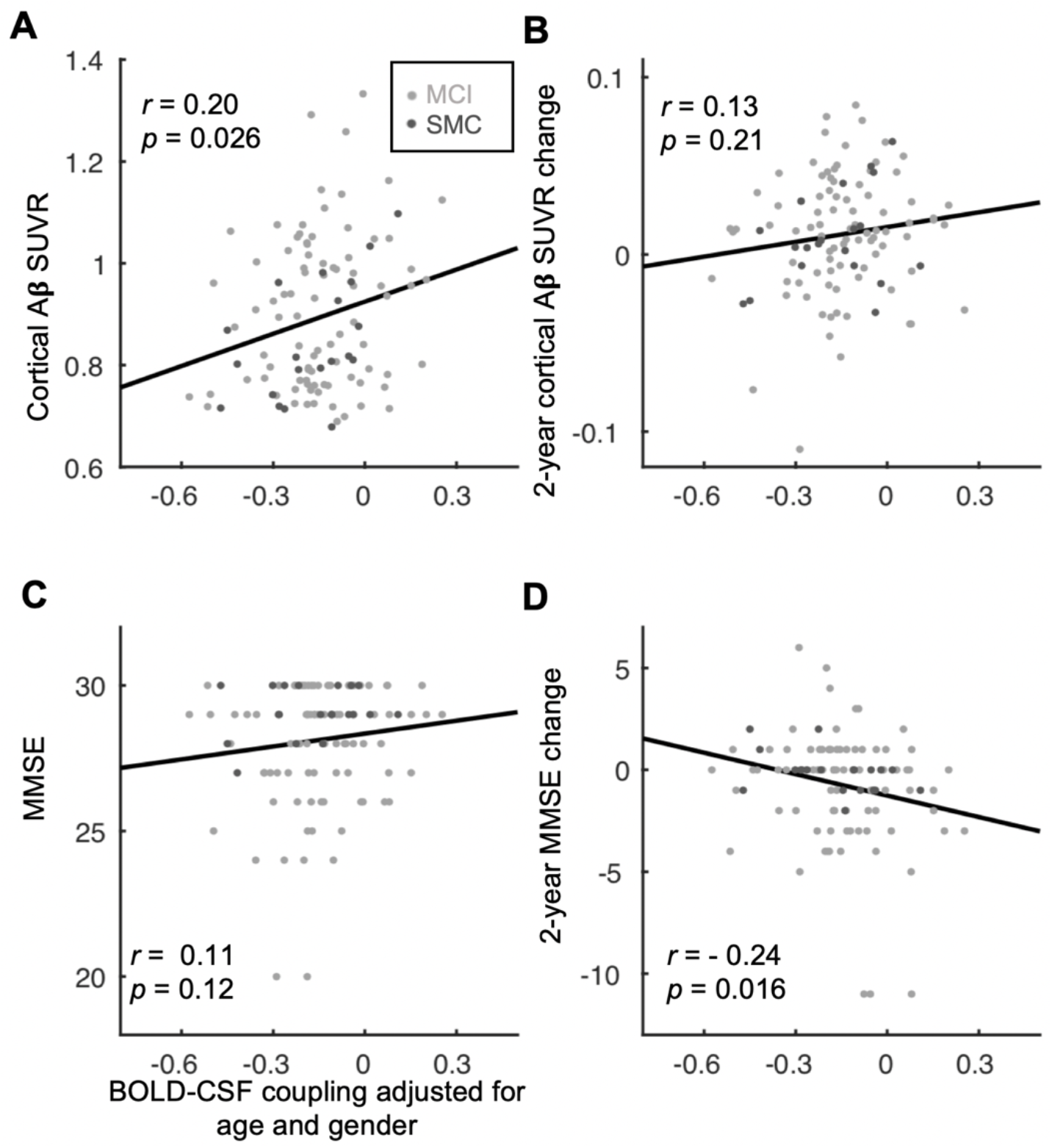
The relationship between AD markers and the BOLD-CSF coupling with excluding the HC and AD data. (**A-B**) The BOLD-CSF coupling adjusted for age and gender is significantly correlated (Spearman’s *r* = 0.20, *p* = 0.026) with the gray-matter Aβ SUVRs (**A**) but not the 2-year longitudinal Aβ changes (**B**) across sessions of the MCI and SMC groups. (**CD**) The BOLD-CSF coupling adjusted for age and gender is significantly correlated with (Spearman’s *r* = −0.24, *p* = 0.016) the 2-year longitudinal changes of MMSE scores (**D**) but not the baseline values (**C**) across sessions from MCI and SMC. Each dot represents a session. MCI and SMC sessions are colored with light gray and dark gray, respectively.

**Fig. S5.**
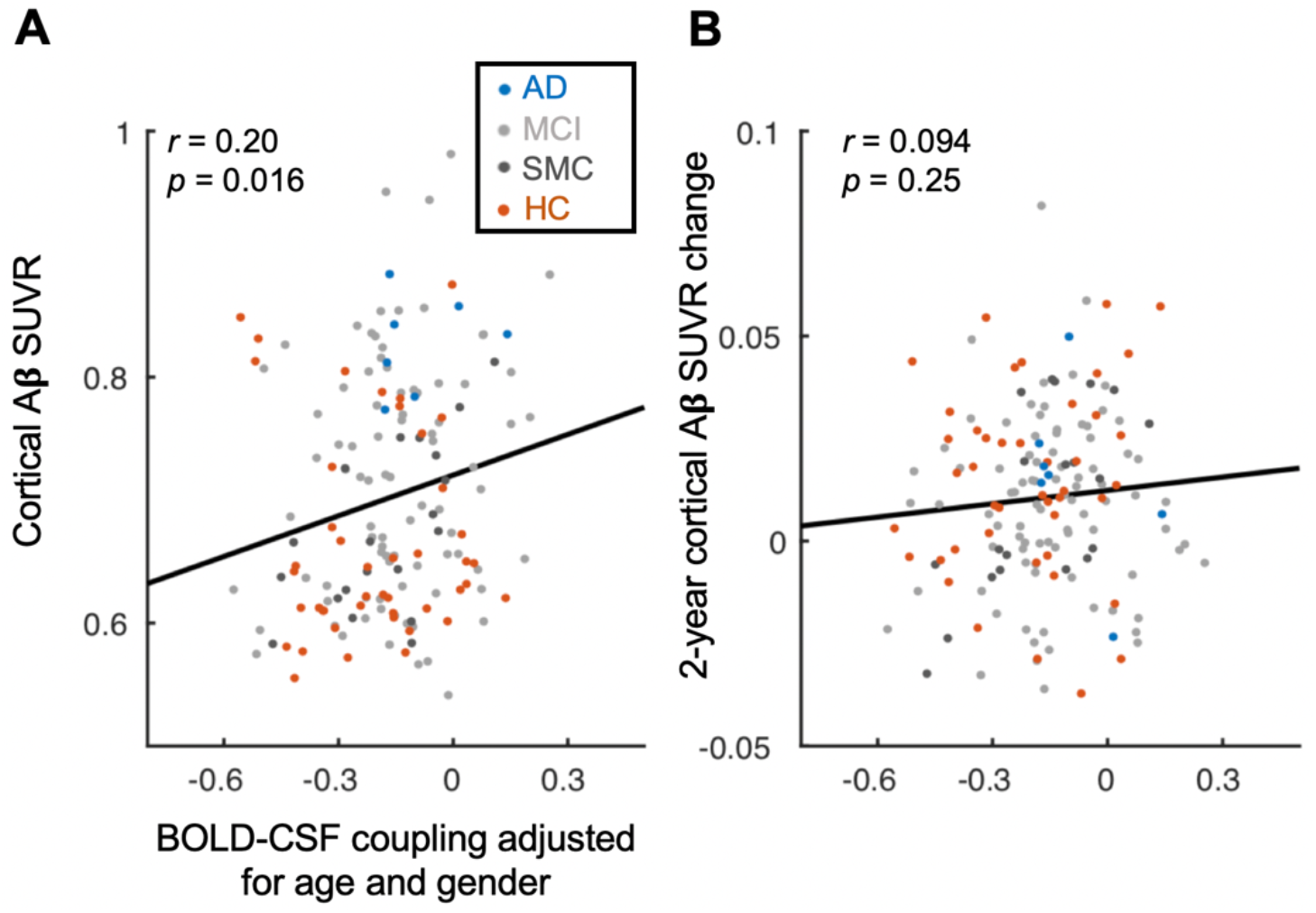
The association between the BOLD-CSF coupling and Aβ-PET SUVR normalized with eroded white matter. (**A-B**) The BOLD-CSF coupling adjusted for age and gender is significantly correlated (Spearman’s *r* = 0.20, *p* = 0.016) with the gray-matter SUVRs (**A**) but not the 2-year longitudinal SUVR changes (**B**) when the reference region is defined as the eroded white matter^40^. The results here with the alternative reference region are very similar to the results in **Fig.3 A** and **B**. Each dot represents a session. AD, MCI, SMC, and HC sessions are colored with blue, light gray, dark gray, and orange, respectively.

**Fig. S6.**
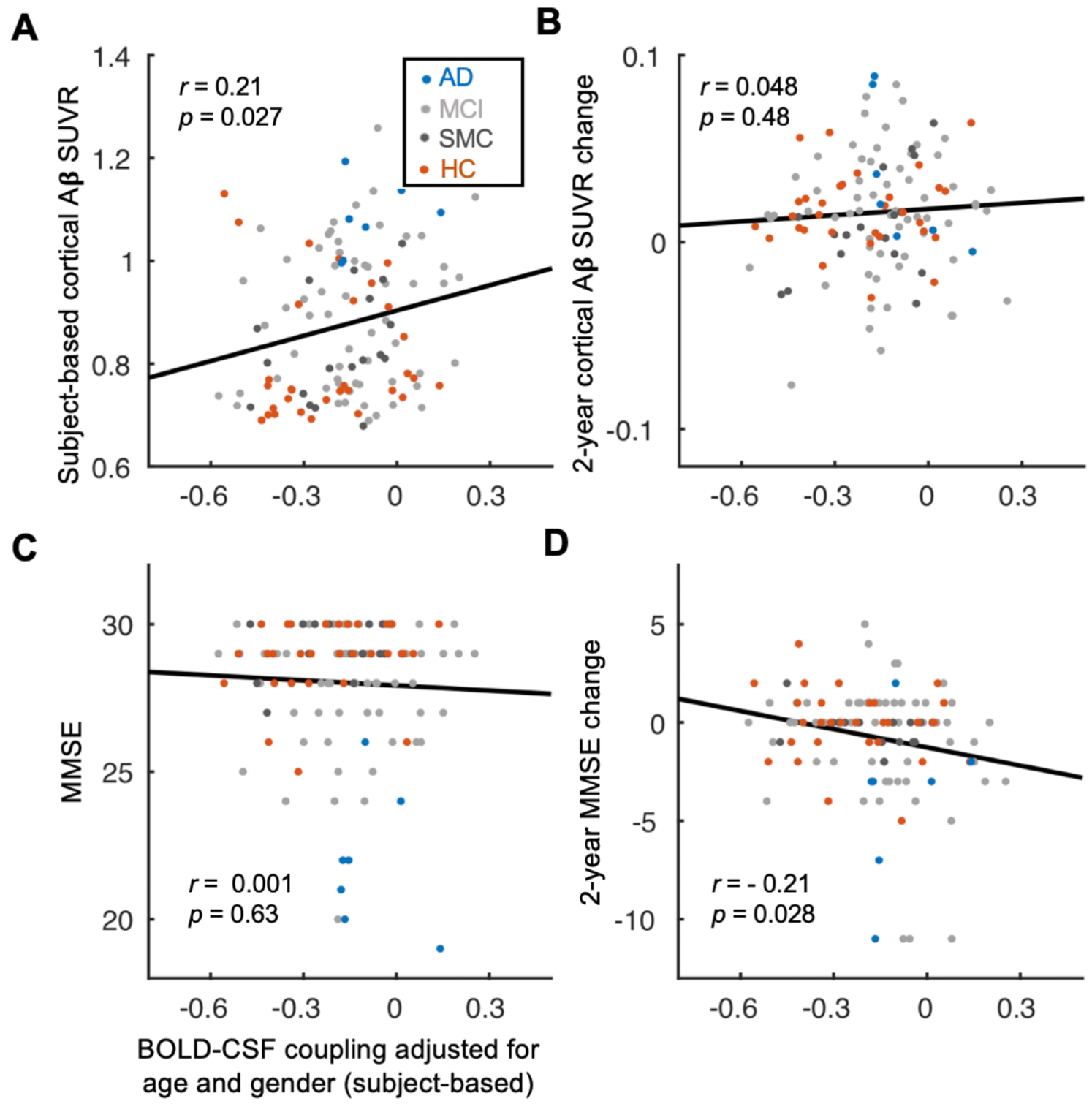
The relationship between AD markers and BOLD-CSF coupling across subjects. (**A-B**) BOLD-CSF coupling adjusted for age and gender is significantly correlated (Spearman’s *r* = 0.21, *p* = 0.027, N = 118) with the gray-matter Aβ SUVRs (**A**) but not the 2-year longitudinal Aβ changes (**B**) across subjects. The composite reference region is the same as **Fig. 3A** and **3B**. (**C-D**) The association between the age- and gender-adjusted BOLD-CSF coupling and the 2-year longitudinal MMSE changes is significant (Spearman’s *r* = −0.21, *p* = 0.028) (**D**) but its correlation with the baseline MMSE score is not significant *(p* = 0.63) (**C**). The linear regression lines were estimated based on the least-squares fitting^78^. Each dot represents a subject. AD, MCI, SMC, and HC subjects are colored with blue, light gray, dark gray, and orange, respectively.

**Fig. S7.**
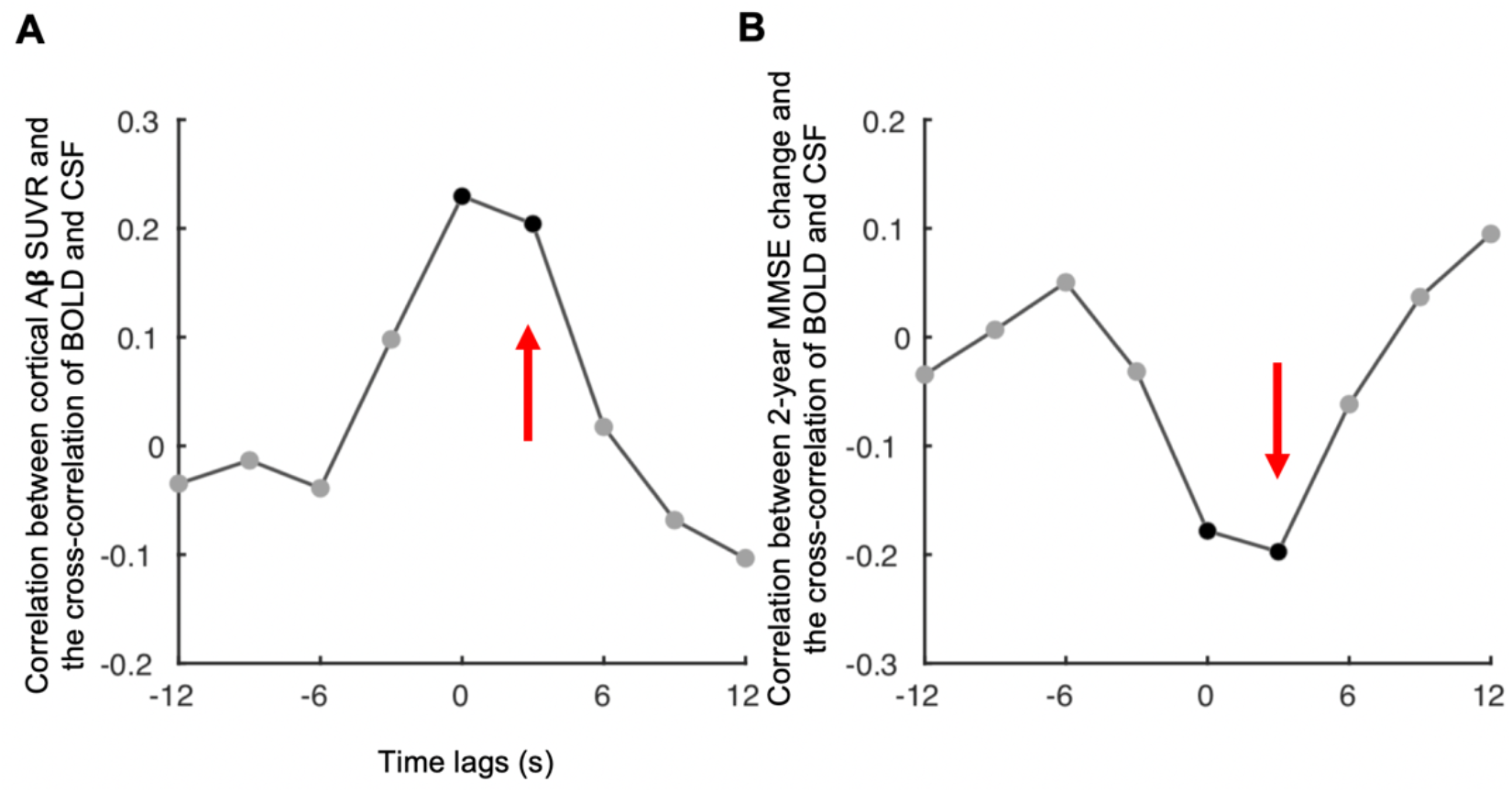
Relationships between AD-related markers and the BOLD-CSF correlations at different time lags. (**A**) The cortical Aβ SUVRs and (**B**) the 2-year longitudinal MMS changes were correlated (Spearman’s *r*, across 158 sessions) with the BOLD-CSF correlations at different time lags after adjusting for age and gender. Black dots indicate significant correlations *(p* < 0.05). Red arrows indicate the lag (+3s) that we used in the “BOLD-CSF coupling” (as **Fig. 2 – 4**).

**Fig. S8.**
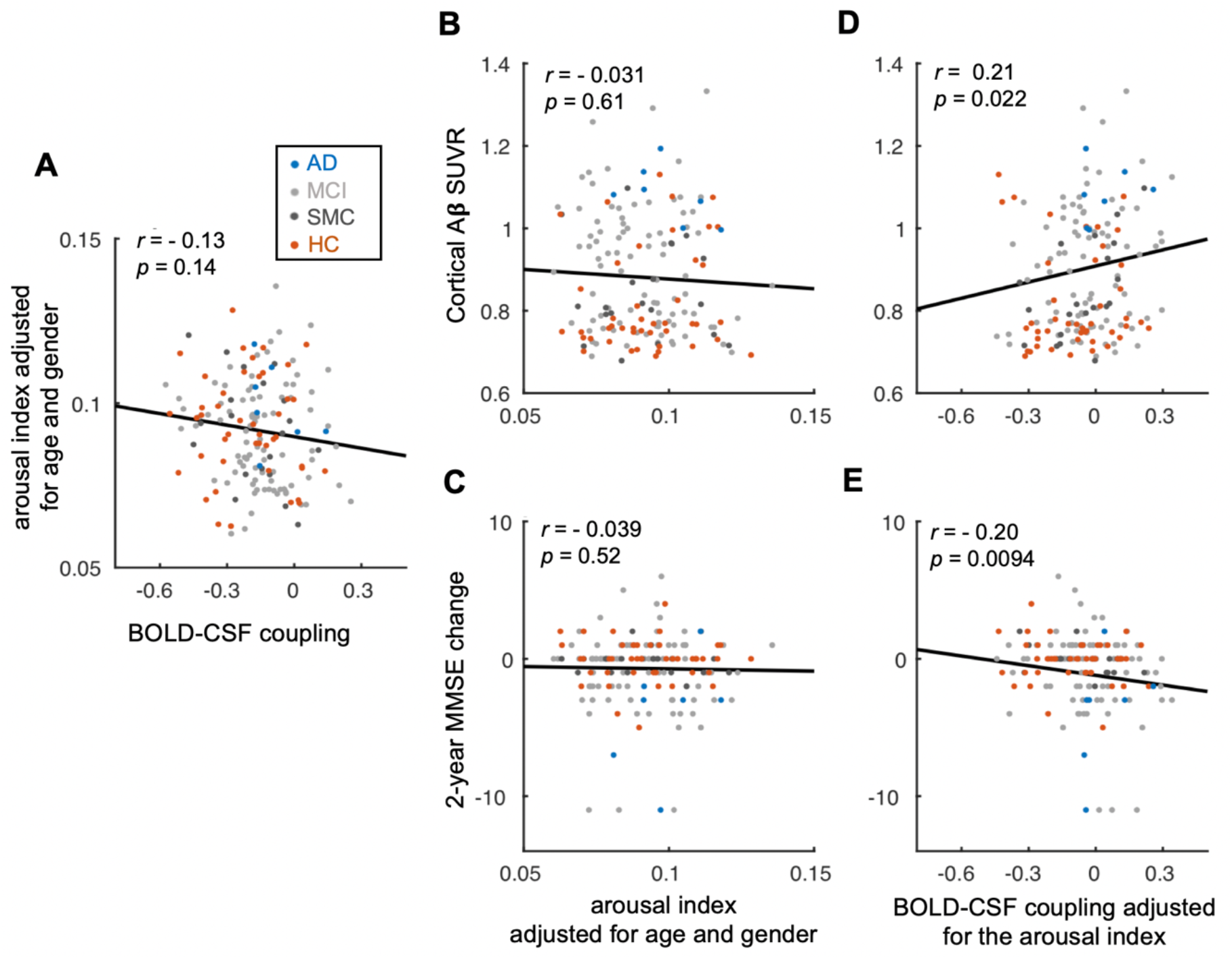
The relationships between the BOLD-CSF coupling and AD-related markers are not affected by the inter-subject variability in arousal. (**A**) The association between the BOLD-CSF coupling (adjusted for age and gender) and the arousal index has a similar trend as **Fig. 4A**. (**B-C**) The arousal index, adjusted for age and gender, is not significantly correlated with either the cortical Aβ level (**B**) or the 2-year longitudinal change of MMSE score (**C**). (**D-E**) The BOLD-CSF coupling remains to be significantly correlated with the cortical Aβ level (**D**) and the 2-year MMSE changes (**E**) after adjusting for age, gender, and arousal index. AD, MCI, SMC, and HC sessions are colored with blue, light gray, dark gray, and orange, respectively. Each dot represents a session.

### Supplementary table

**Table S1.**
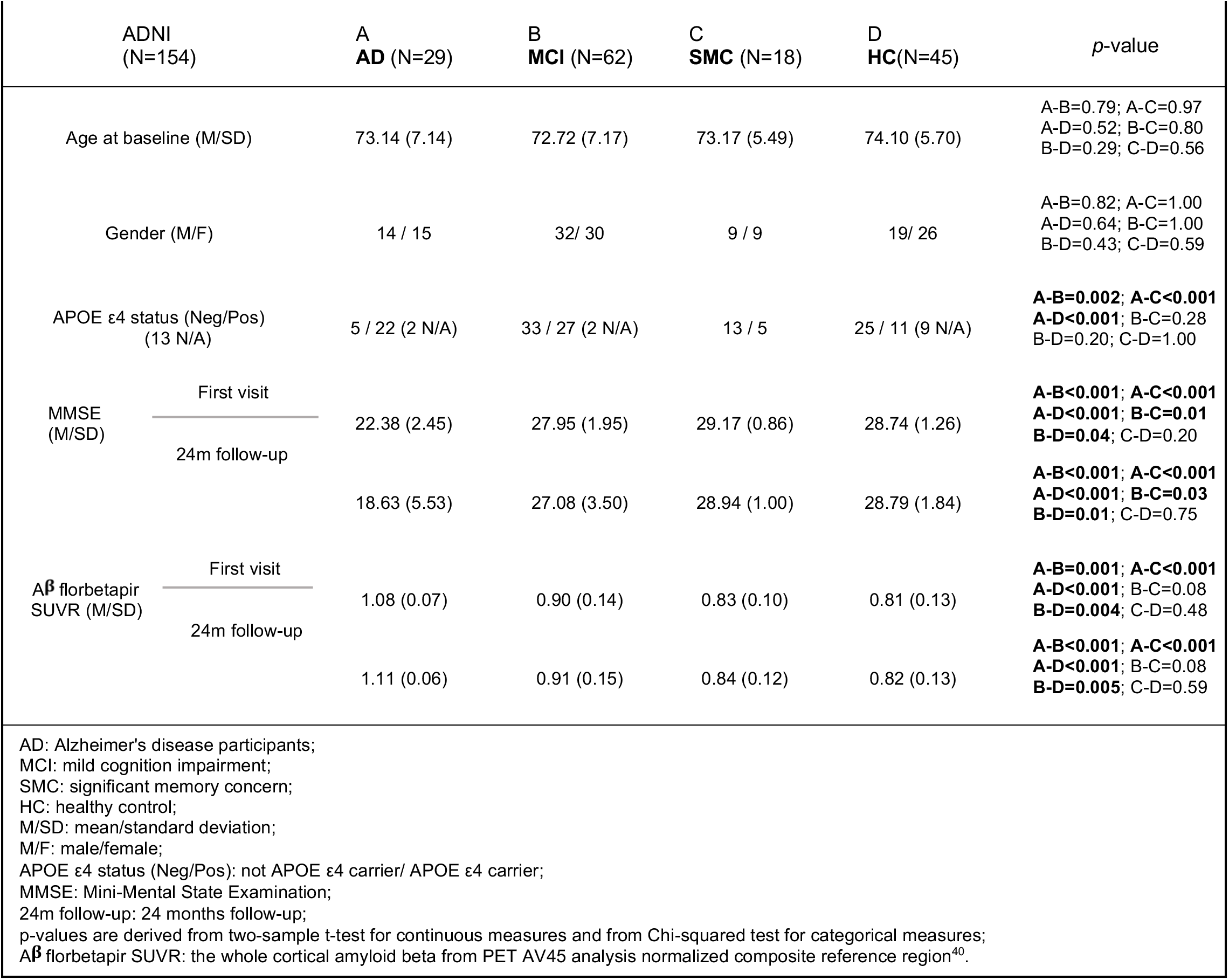
Subject characteristics of the augmented sample.

